# The *Drosophila* pioneer factor Zelda modulates the nuclear microenvironment of a Dorsal target enhancer to potentiate transcriptional output

**DOI:** 10.1101/471029

**Authors:** Shigehiro Yamada, Peter H. Whitney, Shao-Kuei Huang, Elizabeth C. Eck, Hernan G. Garcia, Christine A. Rushlow

## Abstract

Connecting the developmental patterning of tissues to the mechanistic control of RNA polymerase II remains a long term goal of developmental biology. Many key elements have been identified in the establishment of spatial-temporal control of transcription in the early *Drosophila* embryo, a model system for transcriptional regulation. The dorsal/ventral axis of the *Drosophila* embryo is determined by the graded distribution of Dorsal (DL), a homologue of the NF-κB family of transcriptional activators found in humans [1,2]. A second maternally deposited factor, Zelda (ZLD), is uniformly distributed in the embryo and is thought to act as a pioneer factor, increasing enhancer accessibility for transcription factors such as DL [3–9]. Here we utilized the MS2 live imaging system to evaluate the expression of the DL target gene *short gastrulation (sog)* to better understand how a pioneer factor affects the kinetic parameters of transcription. Our experiments indicate that ZLD modifies probability of activation, the timing of this activation, and the rate at which transcription occurs. Our results further show that this effective rate increase is due to an increased accumulation of DL at the site of transcription, suggesting that transcription factor “hubs” induced by ZLD [10] functionally regulate transcription.

## Results

Our study focused on the DL target gene *sog* as its expression domain spans a large dynamic range of the DL gradient, allowing us to examine how ZLD potentiates DL activity across the dorsal/ventral axis. Previous experiments have demonstrated that the lateral stripe of *sog* expression narrows dramatically in *zelda* null embryos [5,11] (Figure 1A,B), and that progressively removing ZLD DNA binding sites from the *sog* shadow enhancer shrinks the domain of activation of reporter genes in a linear manner [7]. In order to understand how ZLD alters transcription at different points along the DL gradient, we revisited these constructs with the aim of visualizing transcription in real time by adding 24 MS2 loops to the 5’ end of the *lacZ* reporter. Since previously utilized MS2 loops [12–15] contained potential ZLD binding sites [16], we revised the MS2v5 [17] sequence to make a ZLD binding site-free non-repetitive version, referred to as MS2v5(-TAG) (see Methods and Table S1). Constructs also contained either the *sog* distal (shadow) enhancer [18,19] with its three native canonical ZLD binding sites, CAGGTAG (hereafter referred to as “3TAG”), or without these sites (hereafter referred to as “0TAG”) (Figure 1C; see Table S2 for enhancer sequences; [7]). The narrowing effect of removing ZLD binding sites was confirmed by *in situ* hybridization (Figure 1D,E).

**Figure 1:**
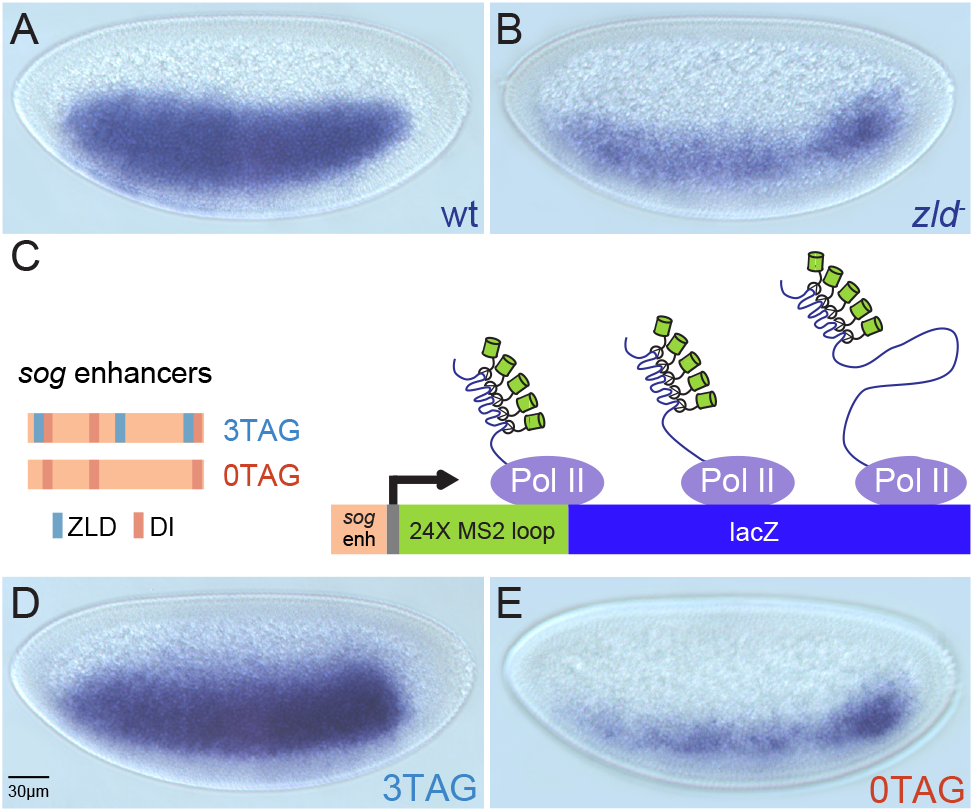
Zelda potentiates Dorsal activity at the *sog* enhancer. (A-B) Conventional enzymatic *in situ* hybridization staining of *sog* in wild type and *zld* mutant NC14 embryos. (C) Schematic representation of transgenes. MS2 loops have been incorporated into the 5’ end of the transcript upstream of a *lacZ* reporter sequence. (D-E) *In situ* hybridization staining for the engineered MS2v5(-TAG) *lacZ* transgenic embryos, showing that 3TAG and 0TAG expression domains phenocopy the expression of *sog* in wild type and *zld* mutants, respectively.

By crossing these transgenic reporter lines to females with a *nanos* promoter-driven MCP-GFP fusion transgene [14], we visualized the transcriptional activation of each reporter as fluorescent foci. These embryos also carry an H2Av(histone 2A variant)-RFP transgene [20], allowing us to track nuclear cycles and record transcriptional activation events in space and time. We performed confocal live imaging over the course of nuclear cycles 10 to 14 (NC10-NC14), tracking the activation of the 3TAG and 0TAG reporter genes (Movies S1 and S2). To validate that the MS2 transgenes behaved as expected, we examined transcriptional activation events in space and time and compared those to expression as assessed by conventional *in situ* analysis. We find that the 3TAG construct is activated as early as NC10, while activation of the 0TAG construct is delayed until NC11-12 (Figure 2A,B; data not shown), in agreement with previously published results of *sog* activation in *zelda* mutants [5].

**Figure 2:**
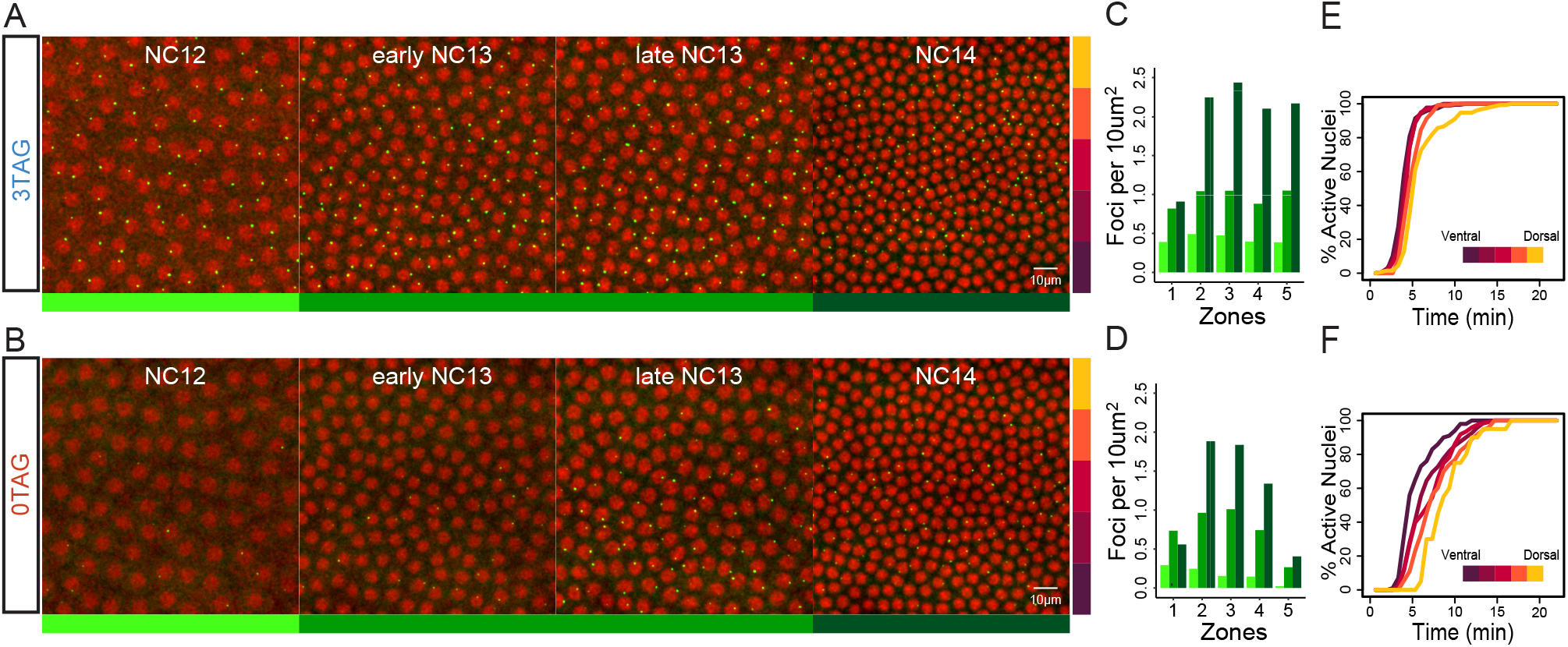
MS2 imaging reveals a position dependent transcriptional delay in the absence of Zelda. (A-B) Frames taken from live imaging movies that track transcription (green spots) from NC12 to NC14 as indicated and color coded below, NC12 (light green), NC13 (medium green), NC14 (dark green). Nuclei (red), have been labeled using maternally loaded H2Av-RFP. Bars on right side represent five zones along the dorsal/ventral axis with ventral mesoderm on bottom (Zone 1). (C-D) Quantification of the number of expressing nuclei in NC 12 to NC14 (color coded as in A-B) agrees with conventional in situ analysis, showing markedly fewer active nuclei in 0TAG embryos across consecutive nuclear cycles, especially in Zones 4 and 5. (E-F) Cumulative distribution curves of nuclei that activate transcription in NC13. 3TAG embryos activate transcription simultaneously across the expression domain, and 0TAG embryos show a delay dependent on the nucleus’ position in the DL gradient (insets are heat maps reflecting Zones 1-5, as in A-B).

To compare the spatial differences in activation, we divided the expression domain of *sog* into five discrete zones with Zone 1 comprising the mesoderm, and all subsequent zones defined by 20μm width bands moving sequentially towards the dorsal midline of the embryo. The *in situ* experiments predict that the most dorsal zones imaged would show few active nuclei in 0TAG embryos, and this was the case. While 3TAG embryos showed similar numbers of active nuclei in each zone across all cycles (NC12-NC14), with the exception of Zone 1 in NC14 due to ventral repression by Snail (Fig. 2C), in 0TAG embryos, the more dorsal the zone, the fewer the number of active nuclei (Fig. 2D). Collectively, these qualitative observations are in accordance with what is currently known about how ZLD participates in transcriptional activation, and provide evidence that our transgenes are faithfully reporting on the transcriptional activity of *sog* in the presence or absence of ZLD.

In addition to allowing qualitative assessment of transcriptional activation, MS2 reporters continually output information on the state of transcription over time, enabling an analysis of the timing of each activation event within a nuclear cycle [14]. This was performed by measuring the time between anaphase of NC12 and the appearance of fluorescent foci in NC13, and plotting the results as cumulative distribution curves (Figure 2E,F). This analysis showed that nuclei in 3TAG embryos express simultaneously across the domain of expression (Figure 2E). In stark contrast, we observed a significant position-dependent delay of activation in 0TAG embryos where the ventral nuclei activate transcription well before lateral nuclei (Movies S1 and S2; Figure 2F). This is presumably due to the highly dynamic nature of the DL gradient, whereby DL levels increase within and across nuclear cycles [21–23]. Here, the 0TAG reporter is effectively acting as a readout for nuclear DL concentration, suggesting that in the absence of ZLD the *sog* enhancer responds to DL levels in a rheostat-like manner, rather than the binary switch-like response seen in the presence of ZLD.

Knowing that activation is altered in 0TAG embryos, we next examined the internal kinetic features of transcriptional activation. We focused principally on two aspects of transcription, diagrammed in Figure 3A. The first was “time to steady-state”, a metric defined as the time between the first instance of signal detection in a given nucleus and the time when that signal reaches maximum output. This is commonly thought of as the time in which a single polymerase molecule has traversed the entire gene body, shown in Figure 3A as *“t”* [14]. The second feature was the “average output at steady-state”, defined by the average signal of the top 20% of values within a given signal trace (see Methods). This variable can be thought of as the average spacing of polymerase molecules on the gene body at max output, expressed as “s”.

**Figure 3:**
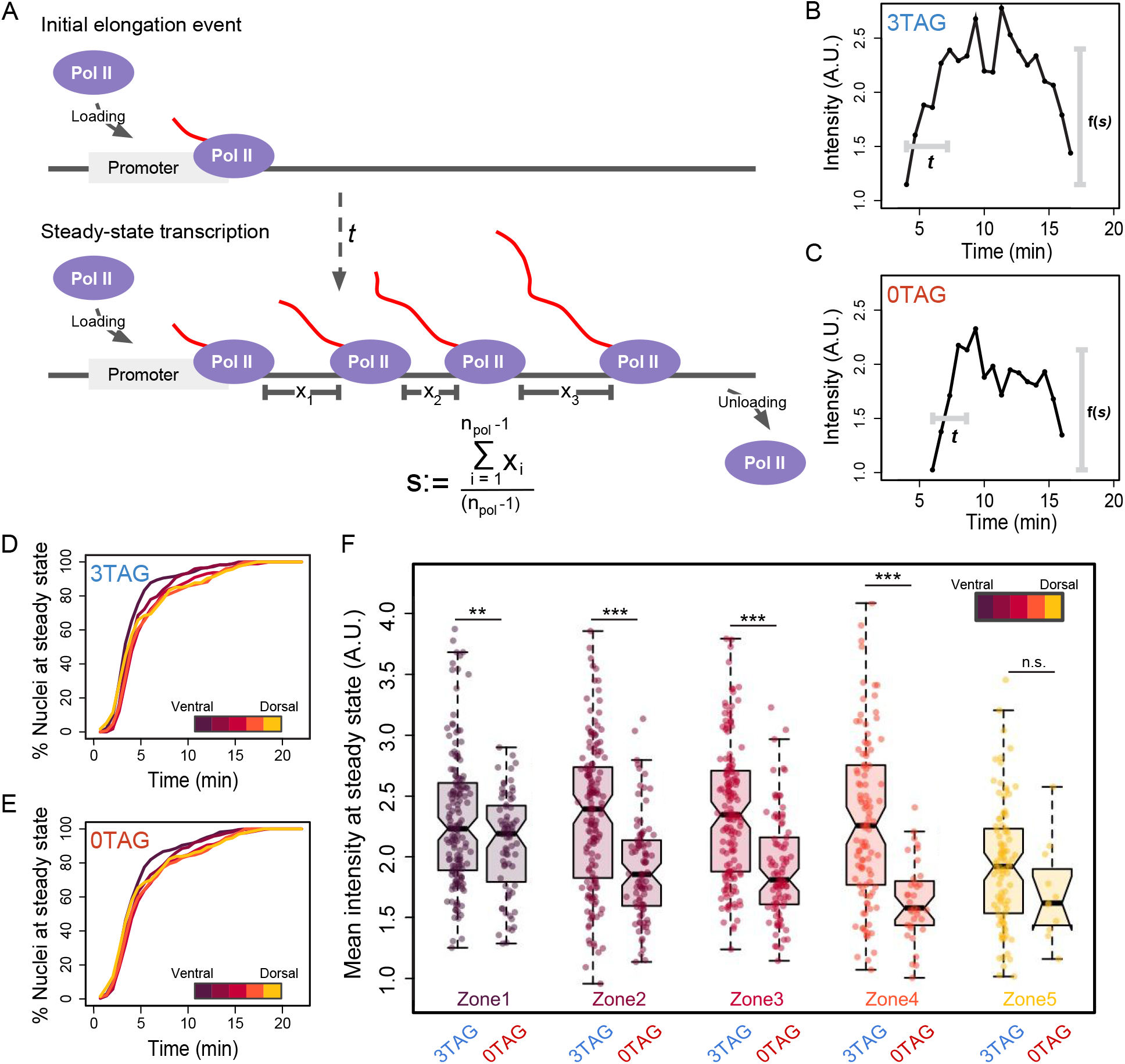
Zelda promotes full saturation of polymerase on the gene body during transcriptional elongation. (A) Schematic representation of the key kinetic steps of transcription following the first productive elongation event. *t* indicates the time to steady-state, the point in transcription where the gene body is decorated with elongating RNA polymerases, and the rate of loading is roughly matched by the rate of unloading. x values show the spacing between polymerase molecules, and s is defined by the average distance between polymerase molecules. This parameter can be modified by multiple biological factors, such as the rate of loading or the speed of escaping a paused polymerase state. (B-C) Example signal traces of single MS2 foci followed through NC13. Parameters from the preceding schematic have been annotated directly onto these traces. *t* is calculated as the length of time between detection above background of the MS2 focus and approximate max output of the gene (see Methods), and max output represented as a function of the average spacing of polymerase molecules. (D-E) Cumulative distribution curves of the percentage of nuclei that have reached steady-state. Neither ZLD nor DL likely play a role in the elongation speed of transcription. (F) Average intensity at steady-state (NC13) plotted as box plot distributions over all 5 zones of the *sog* expression domain. Significant differences between all zones except Zone 5 were found between the genotypes. 3TAG embryos show little difference over the first 4 zones, while 0TAG embryos show progressive loss in signal intensity over the dorsal/ventral axis.

In Figure 3B,C, our kinetic variables have been annotated onto single MS2 output traces, where *“t”* is the time to steady-state, and “f(s)” is a function of the spacing of polymerase molecules across the gene body at steady-state transcription. We expressed the “time to steady-state” results in cumulative distribution curves, with the percentage of all active nuclei at steady-state plotted over time (Figure 3D,E). There is a striking similarity between the two genotypes, indicating that ZLD does not act on the speed of polymerase. In addition, the “time to steady-state” is similar in each of the different zones, suggesting that nuclear DL concentration has little influence on polymerase elongation rate. In contrast, the results for the “average output at steady-state” (Figure 3F) shows both ZLD and DL modulating the strength of transcription. Similar to our observations regarding the onset of transcriptional activation, the 3TAG reporter shows comparable max output across multiple zones until the most extreme end of the DL gradient (Zone 5), whereas the 0TAG reporter shows a progressive loss of max output across the entire gradient (Figure 3F), indicating that transcriptional output rate has become a function of nuclear DL concentration. These results suggest ZLD acts upstream of elongation, for example, to either increase RNA polymerase II loading or decrease the length of pausing experienced by a given polymerase molecule. Either of these regulatory steps would affect the mean spacing of polymerase molecules at max output.

This behavior of ZLD inducing uniform transcriptional activation and output across a transcriptional activator gradient may be explained by ZLD’s reported ability to promote the formation of transcription factor “hubs” [10]. By raising the local concentration of DL at the site of transcription, ZLD may effectively flatten the gradient of DL experienced by the enhancer, and therefore unify the levels of transcriptional output in regions of low level DL. To test this hypothesis, we used a previously described method to examine transcription factor enrichment at sites of nascent transcript formation in *Drosophila* embryos [24,25]. By costaining fixed embryos with an anti-DL antibody and a single molecule (sm) FISH probe targeting the *lacZ* reporter transcript [26], we could quantify the concentration of DL protein adjacent to foci of transcription. Figure 4A shows the DL gradient at comparable positions in 3TAG and 0TAG embryos. Signal overlap between puncta of DL staining and *lacZ* staining, the presumed site of transcription, can be seen in 3D contour maps where the surface represents the level of DL antibody signal and the site of transcription is mapped onto the texture of the contour. We classified nuclei as either having a High, Mid, or Low level of DL based on binning all nuclei imaged according to their average DL signal intensity, which correspond spatially to Zones 1, 2, and 3 in Figures 2 and 3.

**Figure 4:**
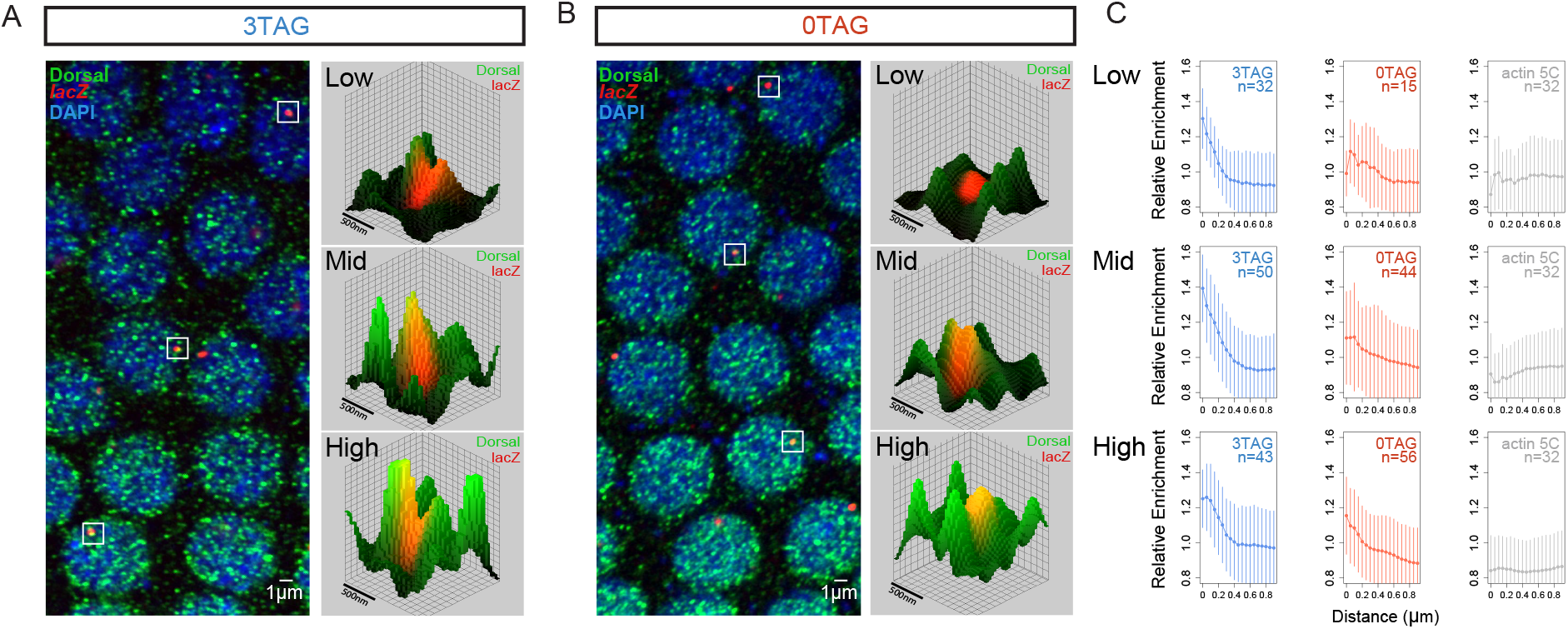
Zelda increases the local concentration of Dorsal at the site of transcription. (A-B) Confocal images of NC13 embryos stained with anti-DL antibodies and smFISH probes for the *lacZ* reporter genes 3TAG (A) and 0TAG (B). DL staining appears highly punctate, indicating the possible presence of high-DL nuclear microenvironments. Sites of active transcription are visualized as red nuclear foci that can be localized in 3D space. Select foci were isolated and visualized in 3D contour maps, where the height of surface represents the intensity of the DL staining. A high incidence of FISH signal overlapping with DL microdomains was observed, suggesting the concentration of DL may have an impact on transcription. (C) The distributions of DL signal within the microdomain of transcribing foci. In regions of high nuclear DL, both genotypes show similar distributions, but a difference is detected in regions where nuclear DL begins to drop. Control distributions were prepared using random places in the nucleus. The numbers of nuclei (n) used for the analysis are indicated. Three embryos for each genotype were used. Error bars: standard deviation.

If transcription is driven by hubs of DL protein, our MS2 experiments predict there should be differences in DL concentration adjacent to the 0TAG enhancer as we move down the DL gradient. Figure 4C uses a modified approach demonstrated by Tsai et al. [27], where the radial intensity of the antibody stain is plotted to visualize the nuclear microenvironment that surrounds a site of active transcription. Because the nuclear concentration of DL changes across the gradient, we divided voxel intensity by the average voxel intensity found within a nucleus. In this way, we could normalize across nuclei by defining our measurement as a unitless index describing the relative enrichment of signal at a given site of transcription, where a value of 1 indicates no enrichment. Additionally, we included a set of random points within nuclei as a control. As predicted, we see a progressive loss in enrichment over the gradient in 0TAG embryos, but no detectable differences in enrichment in 3TAG embryos, indicating that ZLD’s ability to drive higher transcriptional output is based on enhancing the local concentration of existing transcriptional activators rather than utilizing an additional ZLD specific activation pathway. Importantly, these results strongly suggest a functional link between ZLD’s reported ability to induce transcription factor aggregates [10] and transcriptional output, an important first step towards a complete understanding of ZLD’s ability to control gene expression.

## Discussion

Our experiments identify two key parameters where ZLD modifies the activity of a DL-responsive enhancer. The first parameter is the onset of transcription across the domain of *sog*, where a position-dependent delay in transcriptional activation was observed in the absence of ZLD. We believe that the uniformity of this response is the result of ZLD’s pioneering activity to ubiquitously lower the nucleosome barrier from regions of DNA in close proximity to its DNA binding motif. Freeing up enhancers may allow them to be bound more quickly at low concentrations of activators. In the absence of ZLD, DL must compete directly with nucleosomes to access its DNA binding sites. This competition could be more effective at high concentrations of DL, thus leading to the concentration-dependent effects observed in 0TAG.

The second parameter controlled by ZLD is the uniformity of the transcriptional output over the course of a nuclear cycle. Our MS2 data showed remarkably similar levels of total transcription in all measured positions save for the most extreme dorsally-located nuclei. Our results of higher DL enrichment in 3TAG embryos in nuclei with low DL tracks well with the measurements of transcription. DL is an NFκB factor, which activates transcription through the recruitment of the positive elongation factor P-TEFb [28]. A higher concentration of DL would lead to more frequent P-TEFb recruitment, and therefore would increase the rate of polymerase firing provided polymerase loading proceeds independently of any other extrinsic signaling.

The central question our work raises is if these two transcriptional parameters are fundamentally connected by a mechanistic step mediated by ZLD binding to an enhancer. To date, there is no proposed mechanism for how ZLD physically clears nucleosomes. Uncovering the exact mechanism is critical to understanding how transcriptional timing and output are modulated by ZLD. Future experiments are needed to make observations about the dynamics of DL binding. New tools to examine how transcription factors bind to DNA such as single particle tracking will allow visualization of the binding events before and during transcription, and correlate these properties of transcription factor binding directly to transcriptional output.

## Methods

### Drosophila strains

All flies were grown on standard fly cornmeal-molasses-yeast media. *yw* (used as wild type flies), *zld* shmir (*zld*^-^) [8], and transgenic embryos (3TAG and 0TAG) were collected on yeasted grape juice agar plates. Flies of the genotype *y[1] w*; P{His2Av-mRFP1}II.2; P{nos-MCP.EGFP}2* (Bloomington Stock Number 60340) carried two transgenes, one on chromosome 3, <u>*P{nos-MCP.EGFP}2*</u>, which expresses the MS2 coat protein (MCP) fused to EGFP under the control of *nanos*, and the other on chromosome 2, *P{His2Av-mRFP1}II.2*, which expresses RFP-tagged His2Av in all cells under the control of *His2Av*. MS2 transgenes were constructed in the following manner. MS2 loop sequences were revised since previously used MS2 loops [12–14,16,17] contained potential ZLD binding sites [5,14,16]. The new MS2 loops sequence, MS2v5(-TAG) (see Table S1 for DNA sequence) was placed in between the *eve* minimal promoter and a *lacZ* reporter gene (pib-evepr-ms2v5(-TAG)-lacZ plasmid), then subcloned into an attB vector (pBPhi) containing *sog* enhancers with (3TAG) or without (0TAG) ZLD binding sites [7] (Table S1). Transgenic lines carrying these constructs were generated by phiC31 integration in the 53B2 landing site (VK00018), Bloomington stock number 9736 [29,30] by BestGene.

### *in situ* hybridization

Transgenic 3TAG and 0TAG were fixed after bleach dechorionation in 4% formaldehyde (1X PBS) and an equal volume of heptane for 25 minutes while shaking vigorously. Devitellinization was performed by pipetting off the bottom fixative phase and adding 4 ml of methanol and shaking vigorously for 30 seconds. Embryos were rinsed in methanol and transferred to ethanol for storage at −20 degrees C. Hybridization of fixed embryos used a standard in situ hybridization (ISH) protocol and DIG-labeled *sog* or *lacZ* RNA probes and anti-DIG-AP antibodies (Roche Biochemicals) or the Stellaris (LGC Biosearch Technologies) smFISH protocol and Atto-633 conjugated probe sets complementary to *lacZ* (gift from Shawn Little) [26].

### Antibody staining

Antibody staining was performed at 4 degrees for 16 hours followed by three 20 minute washes in PBS + 0.1% Tris pH 7.0. Anti-DL antibody (Dl_7A4) was obtained from the Developmental Studies Hybridoma Bank and used at 1:50 dilution. Embryos were then stained with Alexa Fluor 488 anti-rabbit antibody (Invitrogen, ThermoFisher Scientific) for 1.5 hours and washed in the same manner. After DAPI (D-1388, Sigma-Aldrich) staining for 20 minutes, embryos were mounted on microscope slides using Aqua-Poly/Mount (Polysciences) and Number 1.5 glass coverslips (Fisher Scientific). Embryos were imaged with Zeiss Axiophot DIC optics and a Zeiss Cam and ZEN2012 software. High resolution imaging was performed using a LSM Zeiss 880 confocal microscope with the Airyscan high resolution feature.

### Construction of MS2v5(-TAG) vector

In order to identify potential ZLD binding sites in the DNA sequence encoding MS2v5 [17], the sequence was analyzed with a ZLD alignment matrix (courtesy of Melissa Harrison; [9]) using the Advanced PASTER entry form online (http://stormo.wustl.edu/consensus/cgi-bin/Server/Interface/patser.cgi) [31]. PATSER was run with settings Seq. Alphabet and Normalization “a:t 3 g:c 2” to provide the approximate background frequencies as annotated in Berkeley *Drosophila* Genome Project (BDGP)/Celera Release 1. Complementary sequences were also scored. When PATSER identified a site scoring 3 or higher, one to three bases were modified to reduce the score of the site. After modifying the sequence, it was run through PATSER again to check that no new binding sites were inadvertently created. The process was repeated until all sites scored 3 or higher were abolished. Sites that occurred on sequences encoding MS2 loops were carefully modified to maintain the pattern set forth in Wu *et al*. [17]. Potential binding sites for GAGA Factor were simultaneously abolished during this process using the same methods. The entire MS2v5(-TAG) sequence was constructed as a G-block by GenScript, confirmed by sequencing, and incorporated into our reporter construct by Gibson Assembly (New England Biolabs, Inc.).

### Live imaging

Virgin females maternally expressing MCP-GFP and H2Av-RFP were crossed with males of the MS2 reporter lines. 0-1 hour embryos were collected, dechorionated, and transferred onto a breathable membrane (Lumox Film, Sarstedt AG & Co.; Nümbrecht, Germany) in the middle of a plastic microscope slide (3D printed by Sculpteo; Créteil, France). Live imaging was performed using a Leica SP8 63X objective lens with the following settings: optical sections: 512×512 pixels, 30 z stacks 0.69μm apart, 12bit; zoom: 1.7; time resolution: 40 seconds per frame. Laser power was measured using the X-Cite power meter, model No.XR2100) and set at 70% (main), 30% (488nm), and 10% (554nm). Embryos were imaged for approximately two hours, typically from nc 10 to early nc 14, as *sog* refines rapidly during mid-late nc 14 due to dynamic regulation by other factors [32].

### Quantification and statistical analysis

Live imaging movies were analyzed using the Imaris (Bitplane, Oxford Instruments, Concord MA) “spots” function over and track using retrograde motion with a max frame gap of 3. MS2 foci were assumed to be 1μm across with a z-axis point spread function estimation of 2μm. After tracking, both intensity sum and position csv files were exported and analyzed using a series of custom R scripts. Tracks are assigned a nuclear cycle and zone position by referencing a manually generated annotation file containing all frames where anaphase was reached for each movie and a y-axis position of ventral repression at nuclear cycle 14. Transcriptional delay values for each tracked object are generated by subtracting the current frame number by the preceding anaphase frame number. Transcriptional dynamics at different dorsal-ventral positions was analyzed by subdividing each image into five zones along the DV axis. Zone 1 comprises the mesoderm, as determined by the Snail repression border that becomes obvious in early NC14. The remaining zones are defined by 20μm spatial bins that proceed dorsally, approximately 4 rows of nuclei per zone.

For individual foci traces we performed a linear fit on the first 25% of the intensity values over time. Time to steady-state values were calculated by intersecting the linear fit with a horizontal line generated by the averaging the top 20% intensity values. Statistical tests were performed using Welch’s T-test that assumes independent underlying variance. P-values are visually represented as one asterisk indicating a p < 0.05, two indicating p < 0.01, and three indicating p < 0.001.

smFISH nascent transcript values were obtained by extracting the total fluorescence of large nuclear localized foci assumed to be the point of active transcription. This value was then divided by intensity values of single transcripts by assuming an average 0.3μm diffraction limited point again using the Imaris “spots”. These values formed a normal distribution from which the median value was selected as the fluorescence intensity value of a single transcript within a single frame. DL intensity values for each nucleus were found by extracting the mean fluorescence of antibody stain signal within volumes defined by nuclear DAPI signal. This normalizes differences in DL concentrations along the gradient between genotypes. Radial scans were performed using a custom R script that utilized the position values extracted from Imaris to interrogate .tif files of the DL antibody stain. Error bars on enrichment plots are one standard deviation of all voxels found in each positional bin. All plotting was performed with base R functions and the ggplot2 library.

## Acknowledgements

The authors would like to thank Edo Kussell, Carlos Carmona-Fontaine, Grace Avecilla and Timothee Lionnet for helpful discussions regarding quantitative analysis of MS2 data, and Jonathan Liu, Jacques Bothma, and Stephen Small for critical reading of the manuscript. The research was supported by a National Institute of Health research grant to CAR (GM63024). HGG was supported by the Burroughs Wellcome Fund Career Award at the Scienti1c Interface, the Sloan Research Foundation, the Human Frontiers Science Program, the Searle Scholars Program, the Shurl & Kay Curci Foundation, the Hellman Foundation, the NIH Director’s New Innovator Award (DP2 OD024541-01), and an NSF CAREER Award (1652236).

## Author Contributions

CAR, HGG, and PHW designed the overall study. ECE designed and prepared the MS2v5(-TAG) loops vector. SH made the *sog* enhancer-MS2v5(-TAG) reporter constructs and carried out the live imaging experiments. CAR, PHW and SH carried out the assays in fixed embryos. PHW conceived the ideas and wrote code for the computational image analysis. PHW wrote the manuscript and all authors contributed to revisions.

## Supplemental Information

**Table S1.**
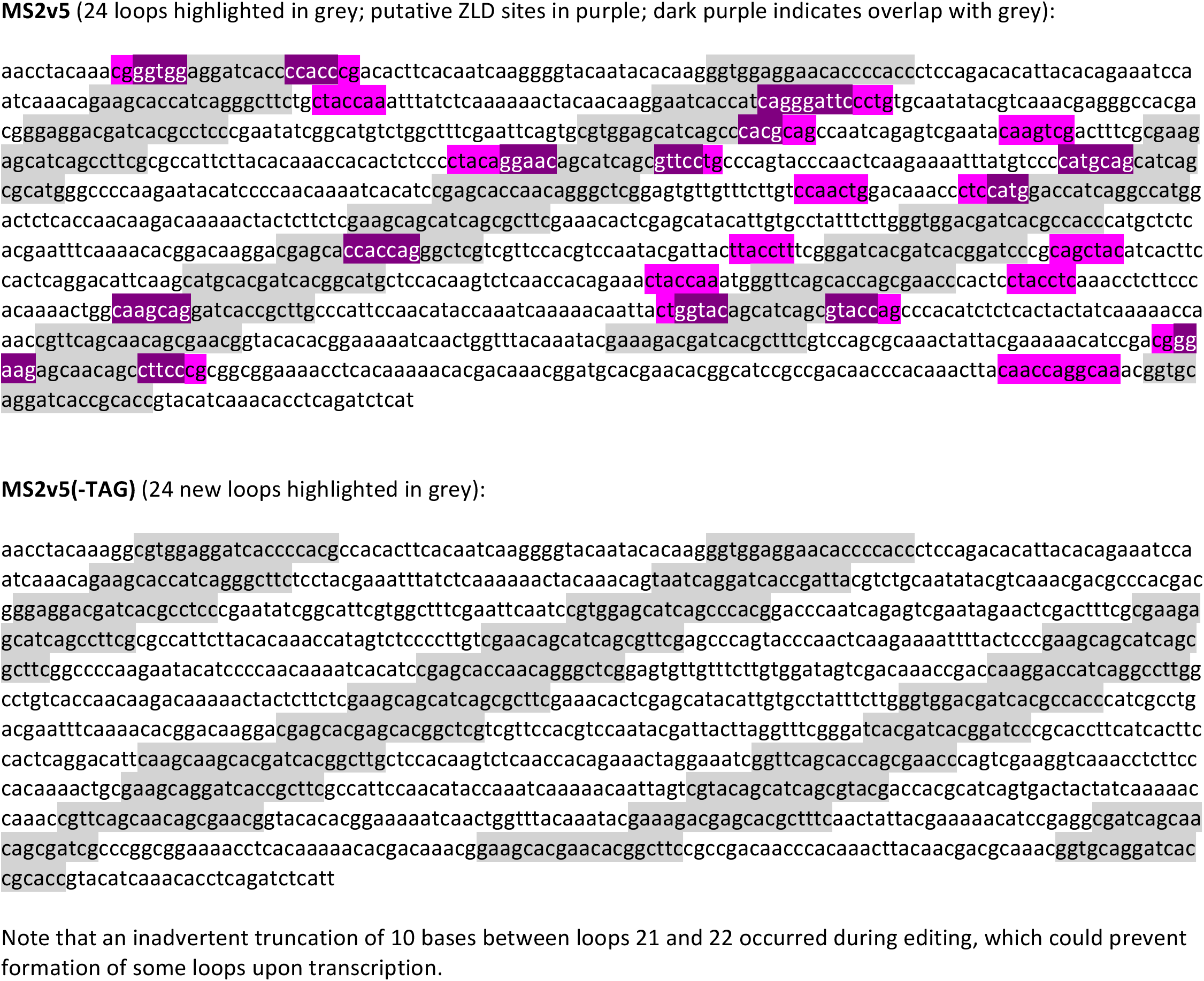

**Table S2.**
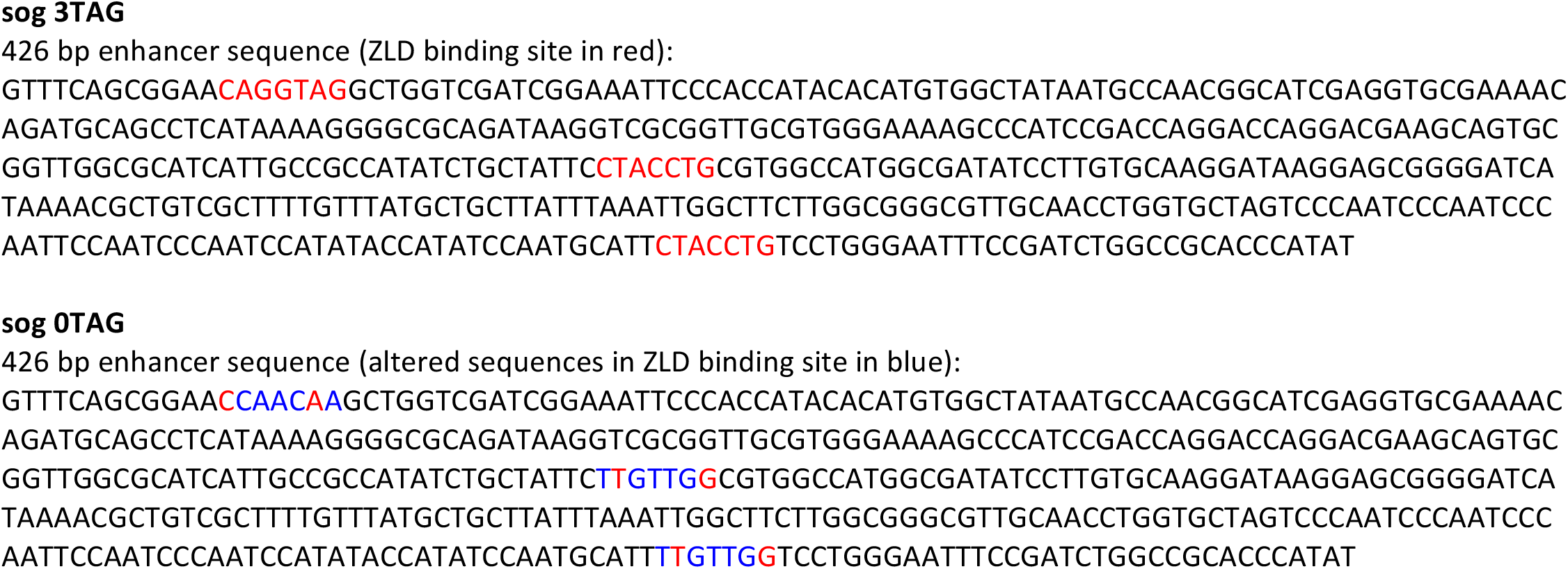

**Movies S1 and S2.**

Time lapse movies of sog 3TAG (051018) (Movie S1) and sog 0TAG (052118) (Movie S2) embryos in NC10-NC14. Embryos were collected from females carrying MCP-GFP (green) and H2Av-RFP (red) mated to males homozygous for the MS2 transgene reporter lines and prepared for live imaging (see Methods) on a Leica SP8 with a 63X objective lens and the following settings: optical sections: 512×512 pixels, 30 z stacks 0.69μm apart, 12bit; zoom: 1.7; time resolution: 40 seconds per frame (see Methods). Scale bar = 10μm

## References

1. Steward, R., McNally, F.J., and Schedl, P. (1984). Isolation of the dorsal locus of Drosophila. Nature 311, 262–265.

2. Stathopoulos, A., and Levine, M. (2002). DL gradient networks in the Drosophila embryo. Dev. Biol. 246, 57–67.

3. Liang, H.-L., Nien, C.-Y., Liu, H.-Y., Metzstein, M.M., Kirov, N., and Rushlow, C. (2008). The zinc-finger protein ZLD is a key activator of the early zygotic genome in Drosophila. Nature 456, 400–403.

4. Schulz, K.N., Bondra, E.R., Moshe, A., Villalta, J.E., Lieb, J.D., Kaplan, T., McKay, D.J., and Harrison, M.M. (2015). ZLD is differentially required for chromatin accessibility, transcription factor binding, and gene expression in the earlyDrosophilaembryo. Genome Res. 25, 1715–1726.

5. Nien, C.-Y., Liang, H.-L., Butcher, S., Sun, Y., Fu, S., Gocha, T., Kirov, N., Manak, J.R., and Rushlow, C. (2011). Temporal coordination of gene networks by ZLD in the early Drosophila embryo. PLoS Genet. 7, e1002339.

6. Harrison, M.M., and Eisen, M.B. (2015). Transcriptional Activation of the Zygotic Genome in Drosophila. Curr. Top. Dev. Biol. 113, 85–112.

7. Foo, S.M., Sun, Y., Lim, B., Ziukaite, R., O’Brien, K., Nien, C.-Y., Kirov, N., Shvartsman, S.Y., and Rushlow, C.A. (2014). ZLD Potentiates Morphogen Activity by Increasing Chromatin Accessibility. Curr. Biol. 24, 1341–1346.

8. Sun, Y., Nien, C.-Y., Chen, K., Liu, H.-Y., Johnston, J., Zeitlinger, J., and Rushlow, C. (2015). ZLD overcomes the high intrinsic nucleosome barrier at enhancers during Drosophila zygotic genome activation. Genome Res. 25, 1703–1714.

9. Harrison, M.M., Li, X.-Y., Kaplan, T., Botchan, M.R., and Eisen, M.B. (2011). ZLD binding in the early Drosophila melanogaster embryo marks regions subsequently activated at the maternal-to-zygotic transition. PLoS Genet. 7, e1002266.

10. Mir, M., Reimer, A., Haines, J.E., Li, X.-Y., Stadler, M., Garcia, H., Eisen, M.B., and Darzacq, X. (2017). Dense Bicoid Hubs Accentuate Binding along the Morphogen Gradient. Genes Dev 31, 1784–1794.

11. Kanodia, J.S., Liang, H.-L., Kim, Y., Lim, B., Zhan, M., Lu, H., Rushlow, C.A., and Shvartsman, S.Y. (2012). Pattern formation by graded and uniform signals in the early Drosophila embryo. Biophys. J. 102, 427–433.

12. Larson, D.R., Zenklusen, D., Wu, B., Chao, J.A., and Singer, R.H. (2011). Real-time observation of transcription initiation and elongation on an endogenous yeast gene. Science 332, 475–478.

13. Forrest, K.M., and Gavis, E.R. (2003). Live imaging of endogenous RNA reveals a diffusion and entrapment mechanism for nanos mRNA localization in Drosophila. Curr. Biol. 13, 1159–1168.

14. Garcia, H.G., Tikhonov, M., Lin, A., and Gregor, T. (2013). Quantitative imaging of transcription in living Drosophila embryos links polymerase activity to patterning. Curr. Biol. 23, 2140–2145.

15. Lucas, T., Ferraro, T., Roelens, B., De Las Heras Chanes, J., Walczak, A.M., Coppey, M., and Dostatni, N. (2013). Live Imaging of Bicoid-Dependent Transcription in Drosophila Embryos. Curr. Biol. 23, 2135–2139.

16. Lucas, T., Tran, H., Romero, C.A.P., Guillou, A., Fradin, C., Coppey, M., Walczak, A.M., and Dostatni, N. (2018). 3 minutes to precisely measure morphogen concentration. Available at: http://dx.doi.org/10.1101/305516.

17. Wu, B., Miskolci, V., Sato, H., Tutucci, E., Kenworthy, C.A., Donnelly, S.K., Yoon, Y.J., Cox, D., Singer, R.H., and Hodgson, L. (2015). Synonymous modification results in high-fidelity gene expression of repetitive protein and nucleotide sequences. Genes Dev. 29, 876–886.

18. Hong, J.-W., Hendrix, D.A., and Levine, M.S. (2008). Shadow enhancers as a source of evolutionary novelty. Science 321, 1314.

19. Ozdemir, A., Ma, L., White, K.P., and Stathopoulos, A. (2014). Su(H)-mediated repression positions gene boundaries along the dorsal-ventral axis of Drosophila embryos. Dev. Cell 31, 100–113.

20. Saint, R., and Clarkson, M. (2000). Pictures in cell biology. A functional marker for Drosophila chromosomes in vivo. Trends Cell Biol. 10, 553.

21. Kanodia, J.S., Rikhy, R., Kim, Y., Lund, V.K., DeLotto, R., Lippincott-Schwartz, J., and Shvartsman, S.Y. (2009). Dynamics of the DL morphogen gradient. Proc. Natl. Acad. Sci. U. S. A. 106, 21707–21712.

22. Liberman, L.M., Reeves, G.T., and Stathopoulos, A. (2009). Quantitative imaging of the DL nuclear gradient reveals limitations to threshold-dependent patterning in Drosophila. Proc. Natl. Acad. Sci. U. S. A. 106, 22317–22322.

23. Reeves, G.T., Trisnadi, N., Truong, T.V., Nahmad, M., Katz, S., and Stathopoulos, A. (2012). DL-ventral gene expression in the Drosophila embryo reflects the dynamics and precision of the dorsal nuclear gradient. Dev. Cell 22, 544–557.

24. He, F., Ren, J., Wang, W., and Ma, J. (2011). A Multiscale Investigation of Bicoid-Dependent Transcriptional Events in Drosophila Embryos. PLoS One 6, e19122.

25. Xu, H., Sepúlveda, L.A., Figard, L., Sokac, A.M., and Golding, I. (2015). Combining protein and mRNA quantification to decipher transcriptional regulation. Nat. Methods 12, 739–742.

26. Little, S.C., Tikhonov, M., and Gregor, T. (2013). Precise Developmental Gene Expression Arises from Globally Stochastic Transcriptional Activity. Cell 154, 789–800.

27. Tsai, A., Muthusamy, A.K., Alves, M.R., Lavis, L.D., Singer, R.H., Stern, D.L., and Crocker, J. (2017). Nuclear microenvironments modulate transcription from low-affinity enhancers. Elife 6. Available at: http://dx.doi.org/10.7554/eLife.28975.

28. Barboric, M., Nissen, R.M., Kanazawa, S., Jabrane-Ferrat, N., and Matija Peterlin, B. (2001). NF-κB Binds P-TEFb to Stimulate Transcriptional Elongation by RNA Polymerase II. Mol. Cell 8, 327–337.

29. Groth, A.C., Fish, M., Nusse, R., and Calos, M.P. (2004). Construction of transgenic Drosophila by using the site-specific integrase from phage phiC31. Genetics 166, 1775–1782.

30. Bischof, J., Maeda, R.K., Hediger, M., Karch, F., and Basler, K. (2007). An optimized transgenesis system for Drosophila using germ-line-specific phiC31 integrases. Proc. Natl. Acad. Sci. U. S. A. 104, 3312–3317.

31. Hertz, G.Z., and Stormo, G.D. (1999). Identifying DNA and protein patterns with statistically significant alignments of multiple sequences. Bioinformatics 15, 563–577.

32. Francois, V., Solloway, M., O’Neill, J.W., Emery, J., and Bier, E. (1994). DL-ventral patterning of the Drosophila embryo depends on a putative negative growth factor encoded by the short gastrulation gene. Genes Dev. 8, 2602–2616.

